# A chromosome-scale assembly of *Brassica carinata* (BBCC) accession HC20 containing resistance to multiple pathogens and an early generation assessment of introgressions into *B. juncea* (AABB)

**DOI:** 10.1101/2022.10.13.512038

**Authors:** Kumar Paritosh, Sivasubramanian Rajarammohan, Satish Kumar Yadava, Sarita Sharma, Rashmi Verma, Shikha Mathur, Arundhati Mukhopadhyay, Vibha Gupta, Akshay K Pradhan, Jagreet Kaur, Deepak Pental

## Abstract

*Brassica carinata* (BBCC) commonly referred to as Ethiopian mustard is a natural allotetraploid containing the genomes of *Brassica nigra* (BB) and *Brassica oleracea* (CC). It is an oilseed crop endemic to the Northeastern regions of Africa. Although it is grown in a limited manner, *B. carinata* is of value as it is resistant/highly tolerant to most of the pathogens affecting cultivated Brassica species of the U’s triangle that are grown worldwide as oilseed and vegetable crops. We report a chromosome-scale genome assembly of *B. carinata* accession HC20 using long-read Oxford Nanopore and Illumina sequencing and BioNano optical maps. The assembly has a scaffold N50 of ~39.8 Mb and covers ~1.11 Gb of the genome. We compared the available long-read genome assemblies of the six species of the U’s triangle and found a highly conserved gene number and collinearity suggesting that *B. juncea* (AABB), *B. napus* (AACC), and *B. carinata* are strict allopolyploids. We cataloged the nucleotide-binding and leucine-rich repeat immune receptor (NLR) repertoire of *B. carinata* resulting in the identification of 465 NLRs. We investigated the extent and nature of early generation genomic interactions between the subgenomes of *B. carinata* and *B. juncea* in interspecific crosses between the two species. We found that C chromosome additions are well tolerated, with homoeologous exchanges occurring between the A and C genomes. Based on the genomic interactions, we propose strategies to utilize the interspecific crosses for transferring disease resistance from *B. carinata* to *B. juncea* and other Brassica species.

## Introduction

The cultivated Brassicas are represented by six interrelated species, three diploids - *Brassica rapa* (2n = 20, genome AA, represented as BraA), *B. nigra* (2n = 16, BB, BniB), and *B. oleracea* (2n = 18, CC, BolC), and three of their natural allotetraploids - *B. juncea* (2n = 36, AABB, BjuA/BjuB), *B. napus* (2n = 38, AACC, BnaA/BnaC), and *B. carinata* (2n = 34, BBCC, BcaB/BcaC). The relationship among these species was first comprehensively described as the U’s triangle (U, 1935). These species, particularly *B. rapa, B. oleracea, B. napus*, and *B. juncea* constitute some of the most extensively grown oilseed and vegetable crops worldwide. *B. napus* oilseed types, called rapeseed, are grown extensively in temperate regions, whereas oleiferous types of *B. juncea* are grown predominantly in the Indian subcontinent. Due to their economic value, there is keen interest in the breeding, genetics, and genomics of the *Brassica* species.

Highly contiguous genome assemblies based on long-read sequencing on PacBio or Oxford Nanopore platforms supplemented with optical maps and/or chromosome capture (Hi-C) are currently available for all the species of the U’s triangle - *B. rapa* (Belser *et al*., 2018; Zhang *et al*., 2018)*, B. nigra* (Paritosh *et al*., 2020; Perumal *et al*., 2020), *B. oleracea* (Belser *et al*., 2018; Guo *et al*., 2021; Lv *et al*., 2020), *B. juncea* (Kang *et al*., 2021; Paritosh *et al*., 2021), *B. napus* (Chen *et al*., 2021; Lee *et al*., 2020; Rousseau-Gueutin *et al*., 2020; Song *et al*., 2020) and *B. carinata* (Song *et al*., 2021). Both breeding activity and genomic information are sparse in the case of the B genome containing Brassicas that are grown mostly by small farm holders in the developing nations.

*B. carinata* originated in northeastern Africa and has been cultivated as an oilseed and a leafy vegetable crop. The Brassica species viz. *B. rapa, B. oleracea, B. juncea*, and *B. napus* exhibit high morphotype diversity and have been domesticated in various agro-ecologies resulting in significant diversity within the species (Cheng *et al*., 2014; Gómez-Campo and Prakash, 1999). *B. carinata* lacks morphotype diversity and has remained localized in the area of domestication till recently. Studies on the germplasm collections of *B. carinata* have reported low levels of genetic diversity in the gene pool (Raman *et al*., 2017; Teklewold and Becker, 2006; Thakur *et al*., 2019; Warwick *et al*., 2006). The narrow genetic base, late maturity, tall plant stature, high erucic acid content, and low yields make *B. carinata* an ineffective replacement for *B. napus* or *B. juncea* as an oilseed crop.

However, *B. carinata* is highly significant for diversification of the other Brassica species gene pools since it contains resistance/high tolerance to many diseases that currently afflict the more widely cultivated Brassicas. *B. carinata* has been reported to be resistant/tolerant to some of the major diseases of the Brassica crops that include powdery mildew (causal agent - *Erysiphe cruciferarum*) (Bradshaw *et al*., 1989; Gong *et al*., 2020; Tonguç and Griffiths, 2004; Uloth *et al*., 2016b), black rot (*Xanthomonas campestris pv. campestris*) (Sharma *et al*., 2016; Taylor *et al*., 2002; Tonguç *et al*., 2003; Vicente *et al*., 2002), Alternaria leaf blight (*Alternaria brassicae*) (Krishnia *et al*., 2000), white leaf spot (*Neopseudocercosporella capsellae*) (Gunasinghe *et al*., 2016), and Sclerotinia stem rot (*Sclerotinia sclerotiorum*) (Navabi *et al*., 2010b; Uloth *et al*., 2016a). *B. carinata* is also tolerant to abiotic stresses such as drought, and heavy metal stress (Enjalbert *et al*., 2013; Limbalkar *et al*., 2021; Malik, 1990; Pan *et al*., 2012; Wei *et al*., 2016).

Here we describe the genome assembly of *B. carinata* accession HC20, using a combination of long-read Nanopore and Illumina sequencing, and BioNano optical maps. The HC20 accession was chosen as it was observed to be highly tolerant to Sclerotinia stem rot and resistant to white rust, and powdery mildew. However, it was only moderately tolerant to Alternaria leaf blight under field conditions. We annotated the HC20 assembly for proteincoding genes, and repeat elements, and compared it with all the available long-read genome assemblies of the U’s triangle species. We also annotated a near-complete set of NLR genes in *B. carinata* and compared their abundance with that in the other Brassica species. We generated interspecific hybrids between *B. carinata* and *B. juncea* and investigated the early generation genome-level interactions between the constituent genomes of the two species. The information from the pilot study has been used to project strategies for introgressions of disease resistance genes/QTLs from *B. carinata* to disease-prone high-yielding cultivars of *B. juncea*. We also suggest ways in which interspecific crosses can be used to improve the yield of *B. carinata*.

## Results

### Genome sequencing and assembly

High molecular weight DNA isolated from *B. carinata* HC20 leaf tissue was used for short- and long-read sequencing. A combination of long-read nanopore sequencing, Illumina reads-based error correction, scaffolding using optical maps, and a genetic map was used to generate the *de novo* assembly of the *B. carinata* accession HC20. The genome of *B. carinata* (BBCC) was estimated to be ~1138 Mb using the kmer frequency distribution of the Illumina paired-end (PE) reads. (Figure S1). Genome sequencing on the Oxford Nanopore MinION and PromethION platforms yielded 4,323,873 reads with an N50 of 21.82 kb providing ~61x coverage of the genome (Table S1). The raw reads were assembled into 1,259 contigs with an N50 of ~10.3 Mb using the Canu assembler (Table 1). The Nanopore reads were used to polish the contigs using the ‘medaka’ program. Illumina PE reads with a ~40x coverage (Table S2) of the genome were subsequently used for four iterative cycles of Pilon-based assembly correction. The quality of the error-corrected contigs was ascertained after each cycle by the number of corrected indels/SNPs (Figure S2).

**Table 1.** Genome assembly statistics of *B. carinata* accession HC20.

|  |  |  |  |
| --- | --- | --- | --- |
| **Oxford Nanopore** | ✓ | ✓ | ✓ |
| **BioNano** |  | ✓ | ✓ |
| **Linkage Map** |  |  | ✓ |
| ***B. carinata* (BBCC, 2n=34)** |  |  |  |
| **Total assembly size (bp)** | 1,100,790,308 | - | - |
| **Number of contigs** | 1259 | - | - |
| **Longest contig** | 36,642,767 | - | - |
| **N50 contig length** | 10,256,563 | - | - |
| **Number of scaffolds** | - | 61 | - |
| **Total scaffold size (bp)** | - | 1,057,784,680 | - |
| **Longest Scaffolds** | - | 70,959,656 | - |
| **N50 scaffolds length** | - | 39,857,088 | - |
| **Un scaffolded contigs** | - | 856(partial) | - |
| **Number of pseudochromosomes/LGs** | - | - | 17 |
| **Scaffolds assigned to LGs** | - | - | 44 |
| **Contigs assigned to LGs** | - | - | 2 |
| **Unassigned scaffolds to LGs** | - | - | 17 |
| **Unassigned contigs to LGs** | - | - | - |
| **Length of assigned sequences to LGs (bp)** | - | - | 1,039,707,227 |
| **Length of unassigned sequences to LGs (bp)** | - | - | 17,824,498 |
| **N50 length pseudochromosomes (bp)** | - | - | 62,332,559 |

Optical mapping was carried out by DLS labeling using the DLE-1 enzyme; a total of 75 Bionano genome maps with an N50 of ~275 Kb were generated (Table S3). The optical maps were used to identify any misassemblies in the contigs and to assemble these into scaffolds. A total of 1259 contigs could be assembled into 61 scaffolds with an N50 of ~39.8 Mb, covering ~1,110 Mb of the *B. carinata* genome (Table 1). A total of 856 unmapped sequence fragments with an N50 of ~68 kb, covering ~54.6 Mb of the genome, remained unscaffolded.

An F_1_DH population of 181 DH lines was derived from a cross of two *B. carinata* lines - HC20 and BEC184 and was genotyped to generate a genetic map. A total of 2044 SNPs, which consisted of 1125 GBS-derived SNPs and 919 Illumina chip-based SNPs, could be mapped onto the 17 linkage groups of *B. carinata*. The map spanned a total genetic length of 2104 cM (Table S4). The genetic markers were used to validate the assembly and generate super-scaffolds. A total of 44 out of 61 scaffolds could be assembled on the 17 pseudochromosomes (Figure 1; Table S5). Two pseudochromosomes contained only one super-scaffold, nine pseudochromosomes contained two super-scaffolds, and the rest contained three or more super-scaffolds. The size of the final *B. carinata* genome that could be assigned to the pseudochromosomes was ~1040 Mb (~ 91.4 % of the estimated genome size of 1138 Mb) (Figure 1; Table 1). Eight pseudochromosomes could be assembled with telomeric repeats on both the ends of the chromosomes, whereas six pseudochromosomes had telomeric repeats at the one end only. We have followed the LG nomenclature of *B. oleracea* var capitata (BolC) (Liu *et al*. 2014) for the BcaC chromosomes, and the nomenclature described for *B. nigra* (BniB) Sangam (Paritosh *et al*., 2020) for the BcaB chromosomes (Table S6).

**Figure 1:**
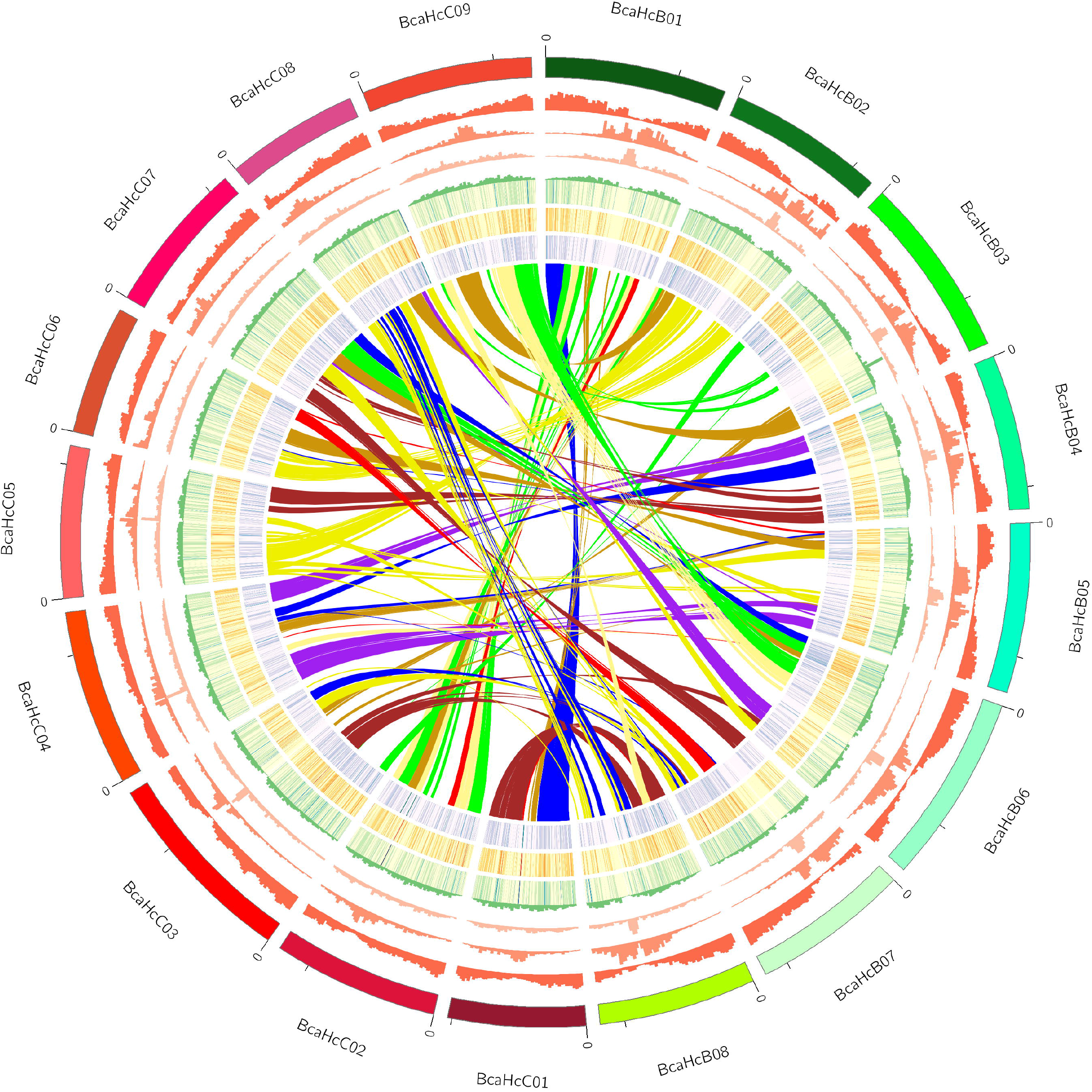
Genomic features of *B. carinata* HC20 assembly. Features (denoted by tracks) on chromosomes are (from outer to inner tracks) (a) gene density per Mb; (b) number of Gypsy elements per Mb; (c) number of Copia elements per Mb; (d) number of LINEs per Mb; (e) number of DNA transposons per Mb; (f) number of simple repeats per Mb; (g) Levels of gene expression in the leaves; (h) Levels of gene expression in the roots; (i) Levels of gene expression in the seeds; (j) Segmental collinearity of the B and C genome components of *B. carinata*. Gene blocks syntenous with *A. thaliana* gene blocks are color-coded following Schranz *et al*. (2006).

To assess the completeness and contiguity of the genome assembly, we used two parameters - a survey of BUSCOs (Benchmarking Universal Single-Copy Orthologs) and the LTR assembly index (LAI). The genome assembly was assessed using three BUSCO databases - Embryophyta, Eudicots, and Viridiplantae (Simão *et al*., 2015). The HC20 assembly had BUSCO scores of 99.6%, 99.8%, and 99.5% against the Embryophyta, Eudicots, and Viridiplantae datasets, respectively (Table S7), indicating that the HC20 assembly was nearly complete for the gene space. The LAI is a reference-free genome metric to assess assembly contiguity by identifying intact LTRs in the assembly (Ou *et al*., 2018). The HC20 assembly had an LAI score of 16.31 (raw LAI score – 18.48), which classifies the HC20 assembly as a “reference-quality” assembly (10≤LAI<20).

### Genome annotation for repeat elements, centromeres, and genes

A *de-novo* prediction approach was used for the identification of transposable elements (TEs). A repeat library developed earlier for the *Brassica* genomes (Paritosh *et al*., 2020; Paritosh *et al*., 2021) was used as a reference. *B. carinata* genome was found to contain ~557.4 Mb (50.13 %) of repeat elements belonging to three broad categories – DNA transposons, retrotransposons, and other repeat elements. The LTR elements were found to be the most predominant repeat elements, constituting ~258 Mb (23.16 %) of the *B. carinata* genome (Table S8). The BcaB genome chromosomes had higher amounts of LTR elements as compared to the BcaC genome (Figure 1). Centromeric repeats were identified in the B and C genomes based on the centromere specific sequences identified earlier (Paritosh *et al*., 2021; Liu *et al*., 2014). For gene annotation, the *B. carinata* chromosome-scale assembly was repeat masked and used for gene prediction with the Augustus program trained with *B. rapa* gene content information as described earlier (Paritosh *et al*., 2020). A total of 1,25,054 proteincoding genes were predicted in the *B. carinata* genome - 60,198 in the BcaB genome and 64,856 in the BcaC genome (Table S9).

We annotated and cataloged some of the important pathway genes in *B. carinata* and compared them to those in the other Brassica species. Specifically, we investigated genes involved in glucosinolate biosynthesis and catabolism, flowering time, and all known transcription factor families. We identified a total of 298 and 342 genes that belonged to the flowering time and glucosinolate pathways, respectively in the *B. carinata* HC20 assembly. We also compared the genes involved in the two pathways across the six species of the U’s triangle (Figures S3-4; Tables S10-11). Additionally, we identified a total of 6246 transcription factors (TFs) belonging to 58 TF families in the *B. carinata* genome (Figure S5; Table S12).

### Transcriptome analysis

RNAseq libraries were generated from nine samples collected from three different tissue types - root, leaf, and 30 days post anthesis seeds of HC20. The libraries were sequenced on the Illumina HiSeq2000 platform; >13 million raw reads were generated for each sample (Table S13). One of the samples (CS2) was removed from further analysis due to low quality of the sequencing data. The cleaned paired-end reads were aligned to the assembled *B. carinata* genome. Pearson correlation between the triplicate samples was >0.95 indicating highly correlated replicates for each tissue type (Figure S6).

A total of 76,486 genes were found to be expressed in the RNASeq of the *B. carinata* HC20 genome. Differential expression analysis identified 14,465, 11,894, and 14,605 genes as upregulated in leaf, root, and seed tissues, respectively (Table S14; Figures S7-9). We further analyzed the leaf, root, and seed-specific genes using the τ tissue specificity score (Yanai *et al*., 2005). A total of 899, 998, and 191 genes were found to be specifically expressed (τ > 0.60) in the seed, root, and leaf tissues, respectively (Table S15). Genes involved in seed maturation [GO:0010431], seed oil body biogenesis [GO:0010344], lipid droplet organization [GO:0034389], seed development [GO:0048316] and lipid transport [GO:0006869] were enriched in the seed transcriptome. Genes involved in the acyl lipid pathway were overexpressed in the seed tissues compared to that in the leaf and root tissues (Figure S9). In contrast, genes related to photosynthesis [GO:0048576, GO:0090229] and phototropism [GO:0010161, GO:0048576] were enriched in the leaf tissues compared to that in the root and seed tissues (Figure S9).

### Gene block arrangement in *B. carinata*

The gene block arrangements of the predicted genes in *B. carinata* were identified by comparisons with the gene blocks marked in the model crucifer *Arabidopsis thaliana* (At) (Schranz *et al*., 2006) using the MCScanX program (Wang *et al*., 2012). Twenty-four gene blocks (A-X) were identified both in the BcaB and BcaC genomes (Figure 2A). Three syntenic regions were identified in the BcaB and the BcaC genomes for each gene block in At as identified in the A, B, and C genomes earlier (Liu *et al*., 2014; Paritosh *et al*., 2020; Wang *et al*., 2011).

**Figure 2:**
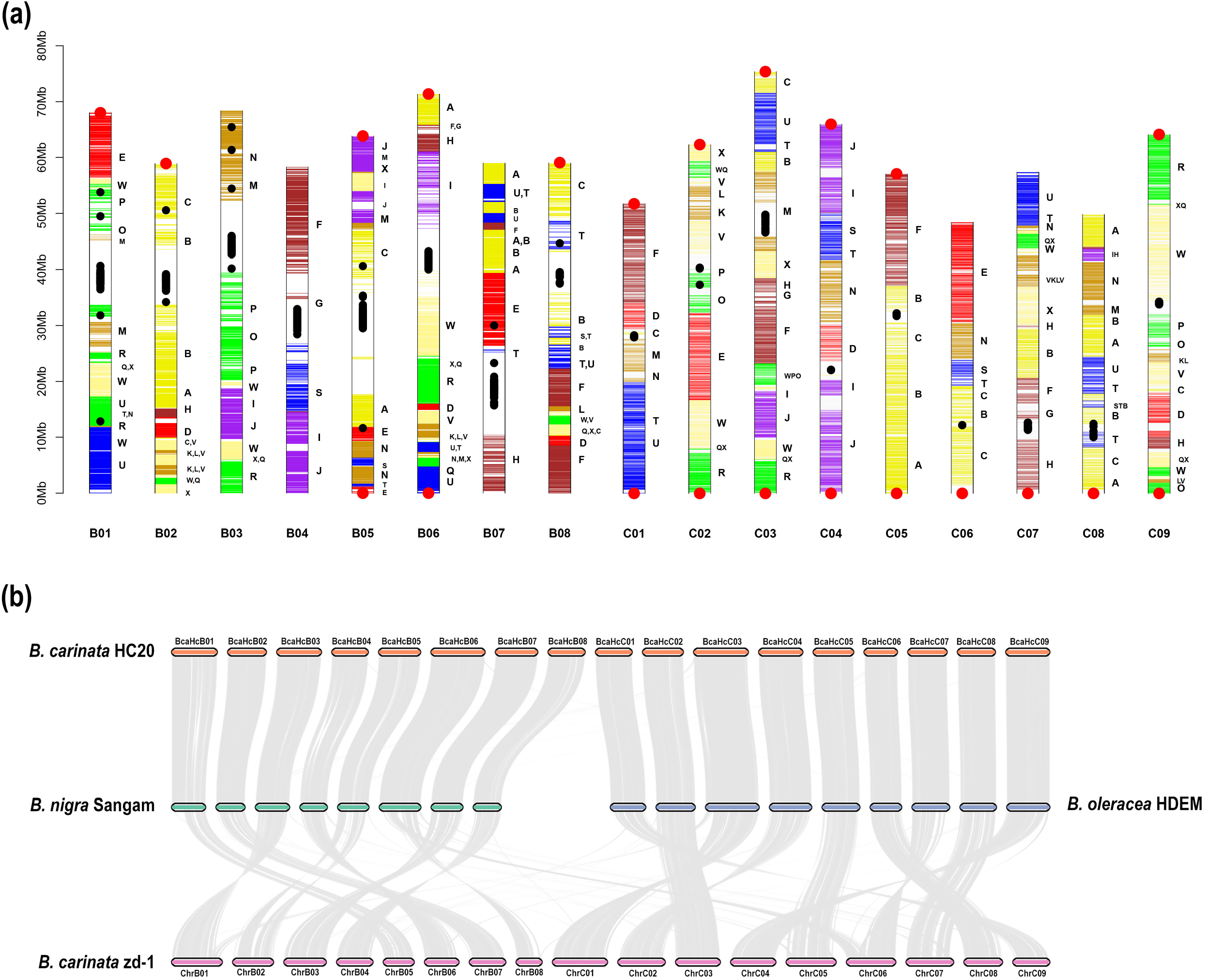
Gene block arrangements on *B. carinata* HC20 pseudochromosomes and extent of synteny with the diploid genomes. (a) Graphical representation of the *B. carinata* HC20 pseudochromosomes. Horizontal bars represent the predicted genes. Different regions of the pseudochromosomes have been assigned to gene blocks based on synteny with *A. thaliana* gene blocks (A–X) as defined by Schranz *et al*. (2006); centromeric and telomeric repeats are represented as black and red dots, respectively. (b) Whole-genome comparison of the HC20 assembly with the diploid genomes *B. nigra* Sangam (Paritosh *et al*., 2020), *B. oleracea* HDEM (Belser *et al*., 2018), and *B. carinata* zd-1 assembly (Song *et al*., 2021).

The data on the physical position and the block assignment of each predicted gene on the 17 *B. carinata* pseudochromosomes BcaHcB01 – BcaHcB08, and BcaHcC01 – BcaHcC09 have been provided in Table S9. The data contains information on the ortholog of each *At* gene in the assembled *B. carinata* constituent genomes – BcaB and BcaC.

### Comparison of the *B. carinata* zd-1 and HC20 genome assemblies

We compared the assembly of *B. carinata* line HC20 with that of *B. carinata* zd-1 published recently (Song *et al*., 2021). We observed large-scale structural differences between the zd-1 and HC20 genome assemblies, especially in the BcaB subgenome. The BcaB genome chromosomes of zd-1 corresponding to BcaHcB01, BcaHcB05, and BcaHcB06 are truncated. Large segmental inversions were present on chromosomes corresponding to BcaHcB03, BcaHcB05, BcaHcC03, and BcaHcC04 (Fig. 2B). Our assembly demonstrated superior quality metrics in terms of BUSCO and LAI scores. The zd-1 assembly had 91.64% complete BUSCO genes (S:22.68%; D:68.96%), while the HC20 assembly has 99.6% complete BUSCO genes (S:5.9%; D: 93.7%). Since *B. carinata* is an allopolyploid genome, the number of duplicated genes is expected to be higher, however, the zd-1 assembly had 24.7% lower duplicated BUSCOs than that in the HC20 assembly. Additionally, the LAI score for the HC20 assembly was 16.31, which was approximately twice that of the zd-1 assembly (LAI: 9.57). The zd-1 assembly is a ‘draft assembly’ as per the LAI classification, whereas the HC20 assembly comes under the category of a ‘reference assembly’.

### Comparative structural genomics of the U’s triangle species

Improved genome assemblies based on the long-read third-generation technologies such as SMRT sequencing on the PacBio platform and Nanopore sequencing have higher levels of contiguity and more extensive coverage of repeat sequences, and gene content (Li *et al*., 2018; Pucker *et al*., 2022). We evaluated a representative genome of each species i.e. *B. rapa* chiifu v3.0 (Zhang *et al*., 2018), *B. nigra* Sangam (Paritosh *et al*., 2020), *B. oleracea* HDEM (Belser *et al*., 2018), *B. juncea* Varuna (Paritosh *et al*., 2021), *B. napus* Darmor-bzh v10 (Rousseau-Gueutin *et al*., 2020), and *B. carinata* HC20 (current assembly) for their gene collinearity on the pseudochromosomes. The A and C subgenomes of the allopolyploids were collinear to the BraA and BolC genomes (Figure 3A). The BjuB and BcaB subgenomes showed high gene collinearity to the BniB genome except for an inversion on BniB04 (Figure 3B). This inversion (in the block J_MF1_-I_MF1_-S_MF2_-S_LF_), however, is specific to *B. nigra* Sangam and has not been observed in the other long-read genome assemblies of *B. nigra* (Perumal *et al*., 2020).

**Figure 3:**
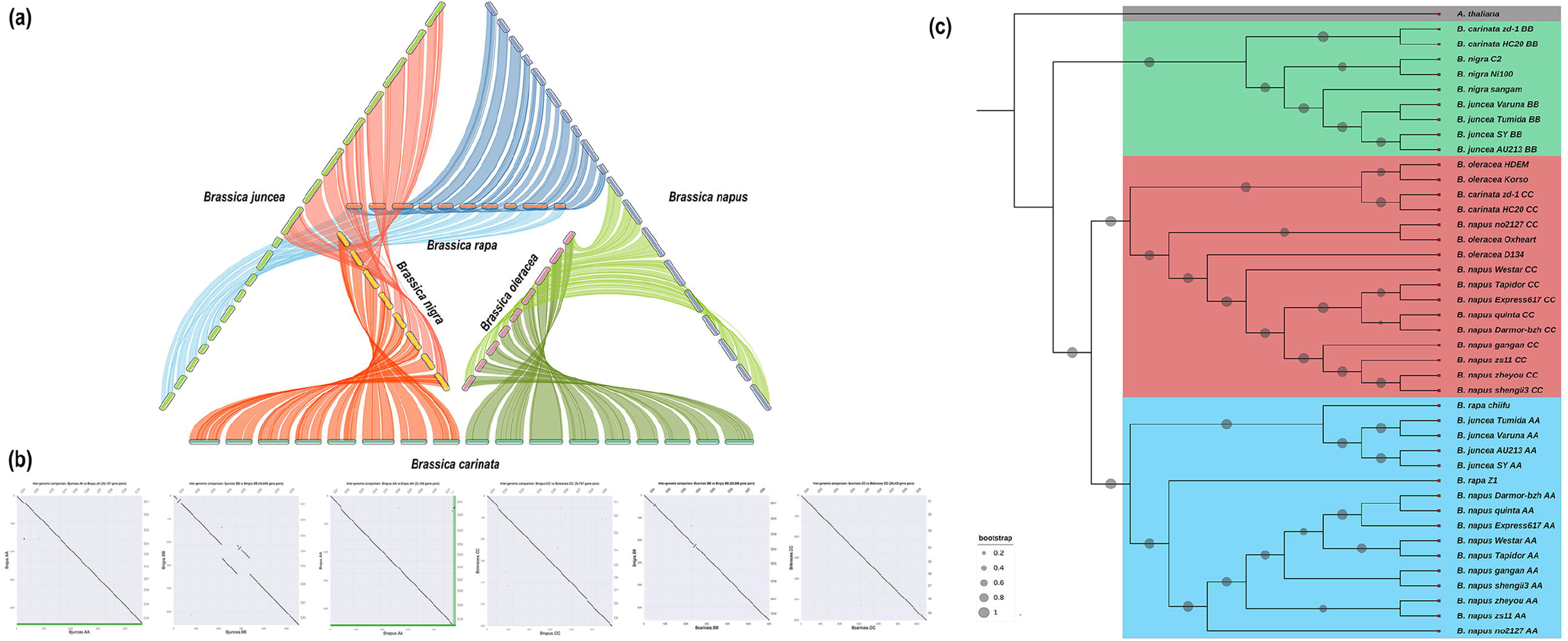
Comparative genomics of the U’s triangle species. (a) Whole-genome comparison of the six *Brassica* species of the U’s Triangle. (b) Dot plots depicting whole-genome comparisons of the allotetraploid genomes with their respective progenitor diploid genomes. The six reference long-read genomes used for the comparisons are *B. rapa* Chiifu v 3.0, *B. nigra* Sangam, *B. oleracea* HDEM, *B. juncea* Varuna, *B. napus* Darmor-bzh v10, *B. carinata* HC20. (c) Phylogenetic tree of the long-read genome assemblies of the *Brassica* species constructed using the 273 single-copy orthogroups. The AA, BB, and CC genomes are colored blue, red, and green, respectively. *A. thaliana*, the outgroup species is colored in grey. Bootstrap values were denoted by circles on the branches, and the size of the circles was proportional to the bootstrap value.

We compared 23 long-read-based genome assemblies (Table S16) of the U’s triangle species published to date to investigate any large-scale structural changes between the BraA, BniB, BolC, and the BjuA/B, BnaA/C, and BcaB/C genome assemblies. We could identify some large-scale inversions in the A and C subgenomes of *B. napus* genotype Express617 (Lee *et al*., 2020) in the BnaA05, BnaA09, BnaC01-04, BnaC07, and BnaC09 chromosomes (Figure S10 & 11). We also observed several structural changes in the BcaB subgenome of *B. carinata* zd-1 (Figure S12) (described in the earlier section). Overall, in all other assemblies, the gene order was maintained in the A, B, and C genomes of the diploid and allopolyploid Brassicas with only minor structural changes.

We also investigated the orthogroups among the six species of the U’s triangle. A total of 69,089 orthogroups were identified, among which 4,663 were species-specific orthogroups. The species-specific orthogroups of most of the species contained < 1% of the total genes included in the analysis (Table S17). The low number of species-specific genes indicates that the gene content is highly conserved among the six interrelated *Brassica* species.

We carried out phylogenetic analysis based on 273 single-copy orthogroups in the 23 long-read *Brassica* genomes. The B genomes clustered distinctly from the A and C genomes (Figure 3C). The analysis revealed that the BcaB genomes were divergent from the BjuB genomes, suggesting evolutionarily distinct B genome ancestors of *B. juncea* and *B. carinata*. This was further corroborated by calculating the mean Ks values of collinear genes between the BniB Sangam genome and the BcaB HC20 and BjuB Varuna genomes. The mean Ks between BcaB and BniB was 0.064, whereas that between BjuB and BniB was 0.059 indicating that the relationship between the BjuB and BniB is closer than that between BcaB and BniB (p-value = 4.688e-10) (Figure S13).

### NLRs of the *B. carinata* and comparison with U’s triangle species

*B. carinata* holds significance for crop improvement of the Brassica species as a donor of disease resistance to many pathogens. Disease resistance can be grouped into (i) qualitative resistance or complete resistance conditioned by a single gene and (ii) quantitative resistance or incomplete resistance conditioned by multiple genes of partial effect (Poland *et al*., 2009). Qualitative resistance is governed by single genes known as nucleotide-binding and leucine-rich repeat immune receptors (NLRs) or R (resistance) – genes. Since we are projecting the use of *B. carinata* HC20 to transfer disease resistance to *B. juncea*, we characterized the whole gamut of NLR genes in the genome. The NLR proteins are classified based on their N-terminal domain into TIR (toll-like interleukin receptor), CC (coiled-coil), and RPW8 (resistance to powdery mildew 8) domain-containing proteins.

We used a combination of the RGAugury pipeline and the NLR-Parser tool to identify NLRs in the *B. carinata* HC20 genome (details in the Materials and methods section). A total of 465 NLRs were identified in the *B. carinata* HC20 assembly, among which 207 and 258 NLRs were located in the BcaB and BcaC genomes, respectively (Figure 4; Table S18). Based on the N-terminal domain classifications, the HC20 genome contained 238 TIR, 72 CC, and 57 -RPW8 domain-containing NLRs. A total of 247 NLRs (53.1%) were grouped in 78 clusters in the HC20 genome. We identified 13% fewer NLR genes in *B. carinata* than in the two diploids combined – BniB Sangam and BolC HDEM. Specifically, the BcaB and BniB genomes had a similar number of NLR genes (BcaB: 203; BniB: 214), whereas BcaC had considerably fewer NLR genes than that present in the BolC genome (BcaC: 257; BolC: 316).

**Figure 4:**
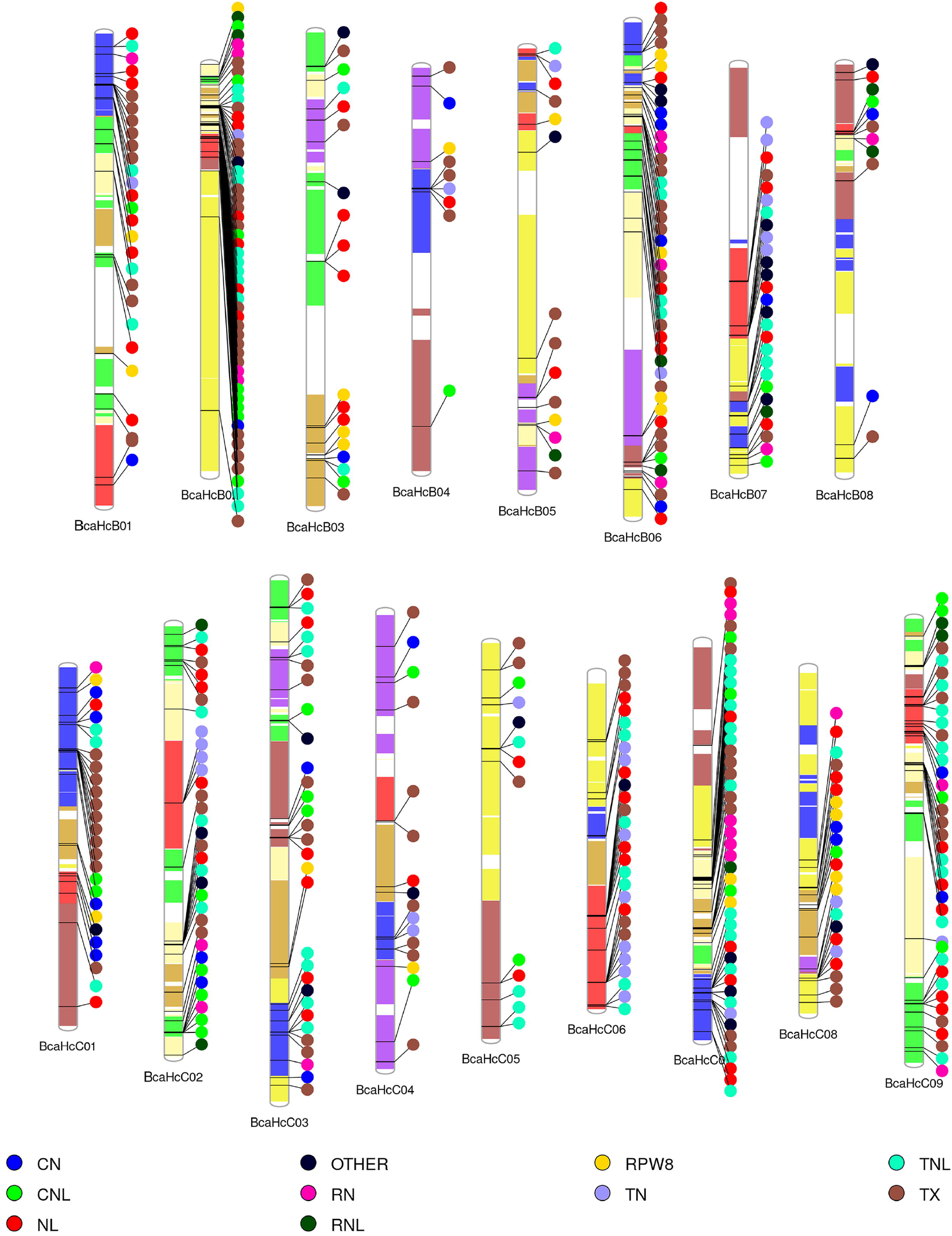
NLR genes in the *B. carinata* HC20 genome. (a) Position of the NLR genes in the 17 chromosomes of *B. carinata*. Different classes of NLRs are denoted by different colors. Chromosomes are colored as per their synteny with the ancestral gene blocks of *A. thaliana*.

The RNLs contain a special N-terminal-resistance domain RPW8 and can confer broad-spectrum resistance against pathogens by themselves or in combination with other domains (Barragan *et al*., 2019). *B. carinata* contains the highest number of RPW8-domain containing NLRs among the Brassica species. The number of N requirement gene 1 (NRG1)-type RNLs are considerably higher in the *B. carinata* genome as 66% (31 of 47 RNLs) of the RNLs clustered in the NRG1 clade compared to that in the ADR1 clade (Figure S14). We identified 48 head-to-head (H2H) NLR gene pairs known as ‘sensor-helper’ or ‘sensor-executor’ pairs in the HC20 genome (Table S19). Thirty of these NLR pairs clustered into two clades corresponding to the sensor-executor NLR pairs identified in *Arabidopsis* (Figure S15) (Van de Weyer *et al*., 2019).

NLRs engaged in the arms race with the pathogenic effectors are fast-evolving genes, which frequently undergo tandem and segmental duplications. Therefore, we carried out a comparative analysis of NLRs in the six species of the U’s triangle. *B. napus* (AACC) had the highest number of NLRs (592) as compared to the other five species (Table S20), which was in concordance with other studies surveying NLR diversity in Brassicaceae (Dolatabadian *et al*., 2020; Inturrisi *et al*., 2020; Tirnaz *et al*., 2020). Most NLR genes are organized into clusters in the six species; however, the composition of the clusters at the syntenic loci varies. Most of the allopolyploid NLRs were syntenic to their parental diploid NLRs, however, *B. juncea* had a higher percentage of syntenic NLRs (78.5%) compared to that present in *B. napus* (64.4%) and *B. carinata* (67.1%) (Figure S16-18). Furthermore, the BcaC and BnaC genomes harbored a higher proportion of non-syntenic NLRs as compared to that in the BcaB and BnaA genomes.

### Early generation assessment of interactions between the BcaBB/CC and BjuAA/BB genomes

Genomic introgressions via interspecific crossing is a powerful approach in crop improvement programs. Introgressions can be limited to introducing select regions or can be large-scale aimed at broadening the genetic diversity of the recipient species. We undertook a study to assess the genomic interactions between the subgenomes of *B. carinata* (BBCC) and *B. juncea* (AABB) to design a strategy for transferring disease resistance from the former to the latter. We used *B. carinata* accessions HC20, HC25, and NPC2 as the male and *B. juncea* Varuna as the female parent as it is a popular variety widely grown in India but is highly susceptible to Sclerotinia stem rot, white rust, Alternaria blight, and powdery mildew.

Besides the HC20 genome assembly reported here, we carried out Illumina short-read sequencing of *B. carinata* accessions used for the crosses viz. HC25, NPC2, and BEC184 at ~10x depth, which revealed the genetic diversity between these accessions (Table S21); the information can be used to develop genetic markers. The F_1_ plants from interspecific crosses viz. VHC1 (Varuna x HC20), VHC2 (Varuna x HC25), and VNP (Varuna x NPC2) were produced by embryo rescue and backcrossed to Varuna as the recurrent parent (Figure S19).

A total of 29 BC_1_F_1_ plants (eight from VHC1, four from VHC2, and 17 from VNP cross, labeled as J28-J56) were grown in the field and genotyped. We used the Brassica 90k Illumina SNP genotyping array to genotype the BC_1_F_1_ plants to track the genomic interactions between the constituent genomes of the two allopolyploids (Table S22). The Brassica 90k array has been earlier used to track the genome-wide changes occurring during interspecific hybridizations (Katche *et al*., 2021; Mason *et al*., 2014a).

The chromosome counts from the array genotyping data in the BC_1_F_1_ plants ranged from 36-45 with a mode of 45 chromosomes (Figure 5; Table S23). The varying number of BcaC chromosomes observed in the BC_1_F_1_ plants indicated that C chromosome additions are well-tolerated. We also observed evidence of homoeologous exchanges (HEs) between the BjuA and BcaC genomes. In the VNP plants, 8.6% of all the BcaC chromosomes exhibited evidence of HEs (Figure 5). In the VHC1 plants, 13.8% of the BcaC chromosomes showed evidence of HEs. The frequency of HEs in the VNP and VHC1 crosses was highest for BcaC01, C02, C04, and C08 chromosomes. The VHC2 plants did not exhibit any evidence of HEs; however, this can be due to the small number of progenies analyzed. One of the progenies (J33) of the VNP cross, resulting from selfing, exhibited extensive genetic exchanges between the BjuA and BcaC subgenomes (Figure 5). However, no viable seeds could be recovered from the field-grown selfed plant.

**Figure 5:**
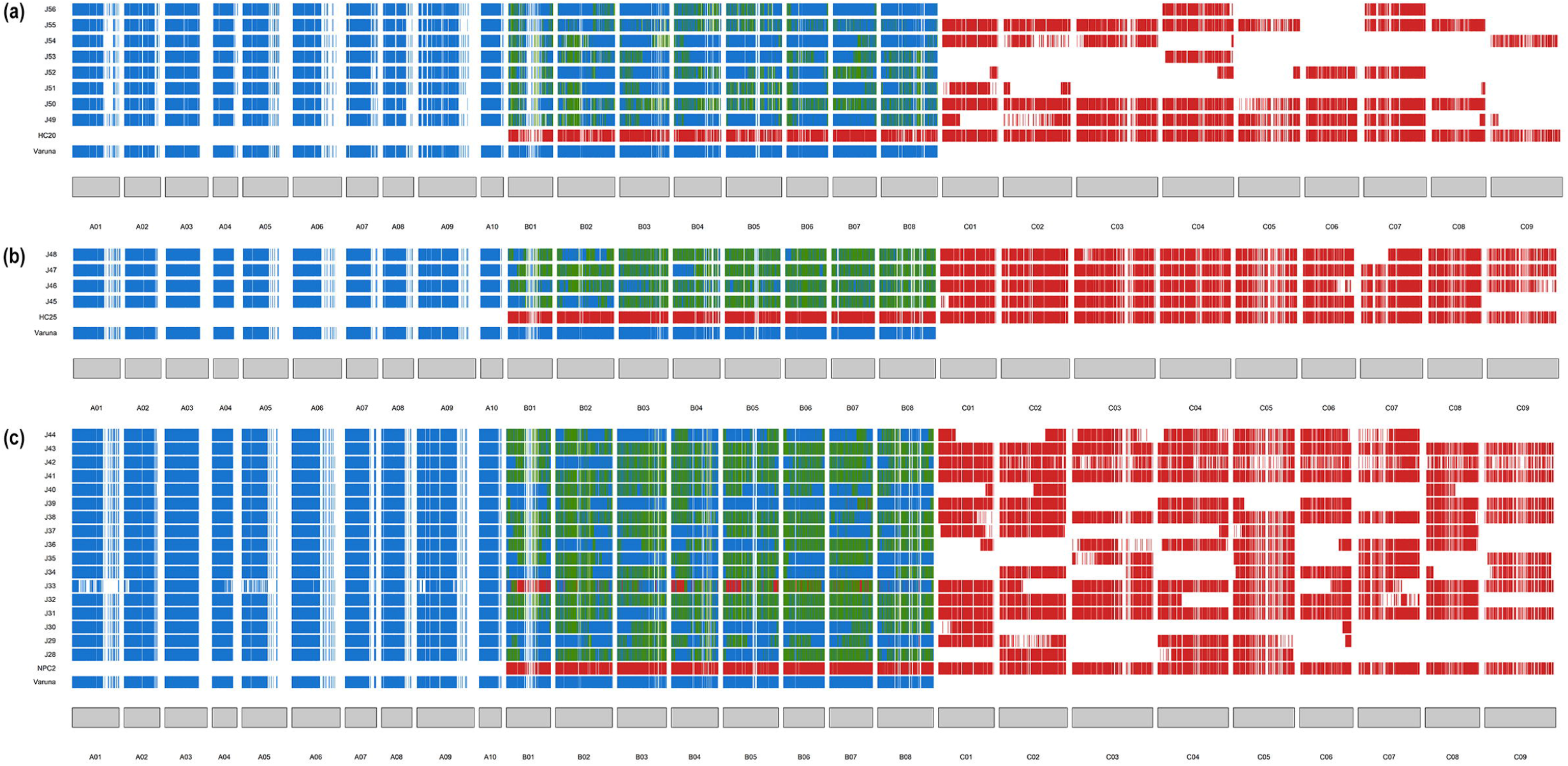
Extent of genomic introgressions from *B. carinata* (BcaB, BcaC) into *B. juncea* (BjuA, BjuB) in the BC_1_F_1_ plants of three independent interspecific crosses VHC1 (a), VHC2 (b), and VNP(c). The blue color represents BjuA-specific SNPs, the red color denotes BcaC-specific SNPs, and the green color denotes BjuB and BcaB heterozygous SNPs. Each colored vertical line across the karyotype represents an SNP. Homoeologous exchanges (HEs) between the BcaC and BjuA chromosomes can be discerned by the presence of BcaC chromosomal fragments without the centromeric region. In all the three crosses, the HEs between BcaC and BjuA mostly took place at the terminal regions. The BcaC chromosome monosomic additions were well tolerated with most lines retaining one or more complete BcaC chromosomes. The line J33 in the VNP cross resulted from selfing in the F_1_ and exhibited extensive rearrangements in the BjuA and BcaC genome chromosomes in addition to some homozygosity for the BcaB genome.

We looked for the stability of HEs in the BC_2_ generation of four BC_1_ plants viz. J29, J44, J40, and J52. The HEs were maintained at reasonably high frequencies in the BC_2_ plants. For example, in the J29-derived BC_2_ progenies, 43.4% of the plants retained the homeologous exchange on the BcaC02 terminal end. Similarly, a HE on BcaC06 was retained in 34.8% of the plants (Figure 6; Figures S20-22). The chromosome additions of the C genome chromosomes were maintained at 20.8 - 91.6 % frequencies in the BC_2_ progenies.

**Figure 6:**
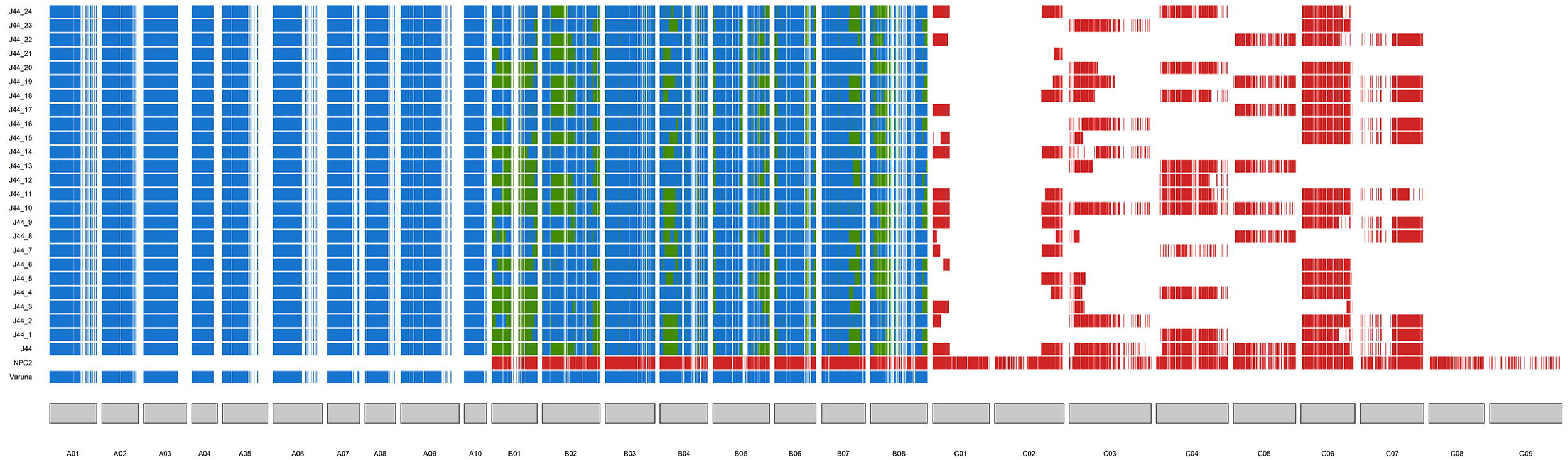
Genomic plots of BC_2_F_1_ lines of J44 BC_1_F_1_ derived from *B. carinata* NPC2 x *B. juncea* Varuna. The three bottom-most lines represent the parental lines *B. juncea* var. Varuna, *B. carinata* NPC2, and J44 (BC_1_ parent). Blue color denotes SNPs of *B. juncea*, red color denotes SNPs of *B. carinata*, and green color denotes heterozygous SNPs. Each line represents an individual with SNPs denoted as colored vertical lines across the karyotype. The BcaC chromosome monosomic additions were retained at high frequencies but segregated in the BC_2_F_1_ progenies. The HEs observed in the BC_1_F_1_ were also retained in the BC_2_F_1_ progenies but underwent recombination in some of the progenies.

Based on the SNP genotyping in the BC_1_F_1_ plants, all 16 BcaB/BjuB chromosomes showed high recombination frequencies but did not recombine with any of the BjuA and BcaC chromosomes. The heterozygosity in the B genome ranged from 7-39%, 15-70%, and 56-67% in the BC_1_F_1_ plants of the VHC1, VNP, and VHC2 crosses, respectively (Table S24). After the second backcross to the recurrent parent *B. juncea* Varuna, the B genome of the BC_2_F_1_ plants showed reduced heterozygosity, ranging from 1.9-52.3% (Figures S20-22; Table S25).

## Discussion

We report a highly accurate and contiguous assembly of the *B. carinata* accession HC20. The genome assembly is a definite improvement over the *B. carinata* zd-1 assembly in terms of gene space and repetitive sequence coverage as evidenced by the assembly metrics such as BUSCO and LAI. The differences in the HC20 and zd-1 assembly could result from two possibilities: 1) the HC20 and zd-1 genomes may have arisen from distinct hybridization events; 2) the differences may be reconciled to the genome assembly artifacts. During the writing of our results, another chromosome-scale assembly of *B. carinata* line Gomenzer was published (Yim *et al*., 2022). This assembly is closer in the quality metrics to the HC20 genome assembly (BUSCOs: zd-1 – 91.6%, Gomenzer – 97%, HC20 – 99.5%; LAI: zd-1 – 9.57, Gomenzer – 12.23, HC20 – 16.31). Additionally, the gene number reported for the Gomenzer assembly (1,33,667 genes) is like that predicted in the HC20 assembly (1,25,054 genes). Although *B. carinata* reportedly has low genetic diversity within the species, whole-genome sequencing of more *B. carinata* accessions is critical to formulating a pangenome for the species that may help elucidate the larger structural variations or presence/absence variations existing within the species.

### Comparative genomic analysis of the U’s triangle species reveals three strict allopolyploids

Our comparative analysis of the U’s triangle species revealed that overall gene collinearity among the six interrelated species was very high with only some minor exceptions (Figures S7-9). The highly conserved gene collinearity indicates that the hybridization of the diploids resulted in strict allopolyploids. In addition, the gene numbers in the polyploids seem to be the sum of the corresponding diploid genomes, indicating that gene retention has been largely favored over gene loss, post-polyploidization (Table S16). We propose that the three allopolyploids are a classical example of sympatric speciation. These allopolyploids in all probability had high initial fertility due to efficient suppression of homoeologous exchanges and heterotic vigor over the co-habiting diploid parents. Polyploid lineages have been reported to undergo either gene retention or extreme gene loss following polyploidization (Bennetzen, 2002; Leitch and Leitch, 2008; Mason and Wendel, 2020). The three Brassica allopolyploids seem to have full retention of the genes of the two parental diploid species.

The meiotic chromosome pairing between homoeologous chromosomes which share a high degree of sequence identity could lead to homoeologous exchanges and has been previously reported in various *B. napus* accessions (Leflon *et al*., 2006; Mason *et al*., 2010; Navabi *et al*., 2010a). Several small-scale HEs have been observed in the natural *B. napus* accessions, whereas large-scale HEs are more prevalent in the synthetic lines (Chalhoub *et al*., 2014; Rousseau-Gueutin *et al*., 2020). However, our analyses of the *B. juncea* (Paritosh *et al*., 2021) and *B. carinata* (current study) genomes suggest that no HEs occur in these genomes. The higher frequency of HEs in *B. napus* may be attributed to the lower levels of divergence between the BnaA and BnaC genomes. However, the Brassica B genome, which diverged earlier from the A and C genome, has a more distinct gene block arrangement compared to that in the A and C genomes (Paritosh *et al*., 2020).

### The NLR repertoire of *B. carinata* and comparisons with other Brassica species

The *B. carinata* HC20 genome contained 465 NLRs – 207 and 258 NLRs in the BcaB and BcaC subgenomes, respectively. The HC20 genome had a total of 153 NLRs that were not present in syntenic positions in the diploid Brassicas (BniB and BolC). The nonsyntenic or novel NLRs in *B. carinata* may possibly contribute to the broad-spectrum resistance against various biotrophic pathogens such as *A. candida* and *E. cruciferarum*.

The variable number of NLR genes predicted in different studies may result from different NLR annotation approaches, quality of genome assembly, and the level of repeat masking. In our comparative analysis, we have used a uniform approach to gene prediction and annotation to avoid these discrepancies. Our analyses of NLR genes in Brassica species revealed that > 50% of them exist as clusters in the genome (Figure 4; Table S18). The clustered arrangement of NLR genes may be vital to the evolution of the NLRs via recombination during meiosis. Recombination between these clusters can change the NLR numbers, induce domain reshuffling, and increase sequence diversity thereby playing a crucial role in the arms race between the pathogen and the host (Ameline-Torregrosa *et al*., 2008; Yu *et al*., 2014; Zhang *et al*., 2016).

Species-specific NLR gene duplications and clustering patterns have been reported earlier (Dolatabadian *et al*., 2020; Inturrisi *et al*., 2020; Zhang *et al*., 2021; Zhang *et al*., 2016). In our study also, we find that NLR clusters in the B-genome of diploids and tetraploids are collinear, however, there are changes in the NLR gene content within these clusters. The presence of NLR genes in clusters could aid in the transfer of these genes between species, as a single functional unit, without disrupting their putative interactions.

### Genomics-assisted introgression of *B. carinata* into *B. juncea* for disease resistance

*B. carinata* (BBCC) is a suitable donor for improving environmental resilience and disease resistance in the high-yielding cultivars of *B. napus* (AACC) and *B. juncea* (AABB) through interspecific hybridizations (Limbalkar *et al*., 2021; Raman *et al*., 2017). The possibility of introgressions is higher in *B. juncea* as it shares the B genome with *B. carinata* and the A and C genomes have a higher propensity for recombination with each other than with the B genome (Attia *et al*., 1987; Mason *et al*., 2010; Struss *et al*., 1991). A recently published study on interspecific crosses between *B. carinata* and *B. juncea* using SNP markers demonstrated perfect pairing between the BcaB and BjuB genomes and recombination and restructuring of the BjuA and BcaC genomes (Katche *et al*., 2021). The selection in each generation of the selfed progeny was for high fertility and genomic stability.

The current study on *B. carinata* and *B. juncea* crosses was aimed at tracking the interactions between the subgenomes for introgressions of disease resistance. The genetic loci conferring resistance to different pathogens in *B. carinata* have not been mapped in crosses within the *B. carinata* germplasm. Therefore, the interspecific hybridization of *B. carinata* and *B. juncea* has two facets – i) the crosses need to be utilized for identifying and mapping the chromosomes/chromosomal regions that confer resistance, and ii) to introgress the identified regions into *B. juncea* for disease resistance. Based on the results of the interspecific crosses carried out in this study, we propose a strategy for genomics-assisted introgression of the *B. carinata* genome into *B. juncea* for disease resistance against major pathogens (Figure 7). The initial crosses of *B. carinata* with *B. juncea* should be followed up by one generation of backcrossing to *B. juncea* since the selfing of the F_1_ hybrid leads to poor fertility and deleterious chromosome configurations. A large number of BC_1_F_1_ progenies need to be generated and analyzed for their chromosomal makeup.

**Figure 7:**
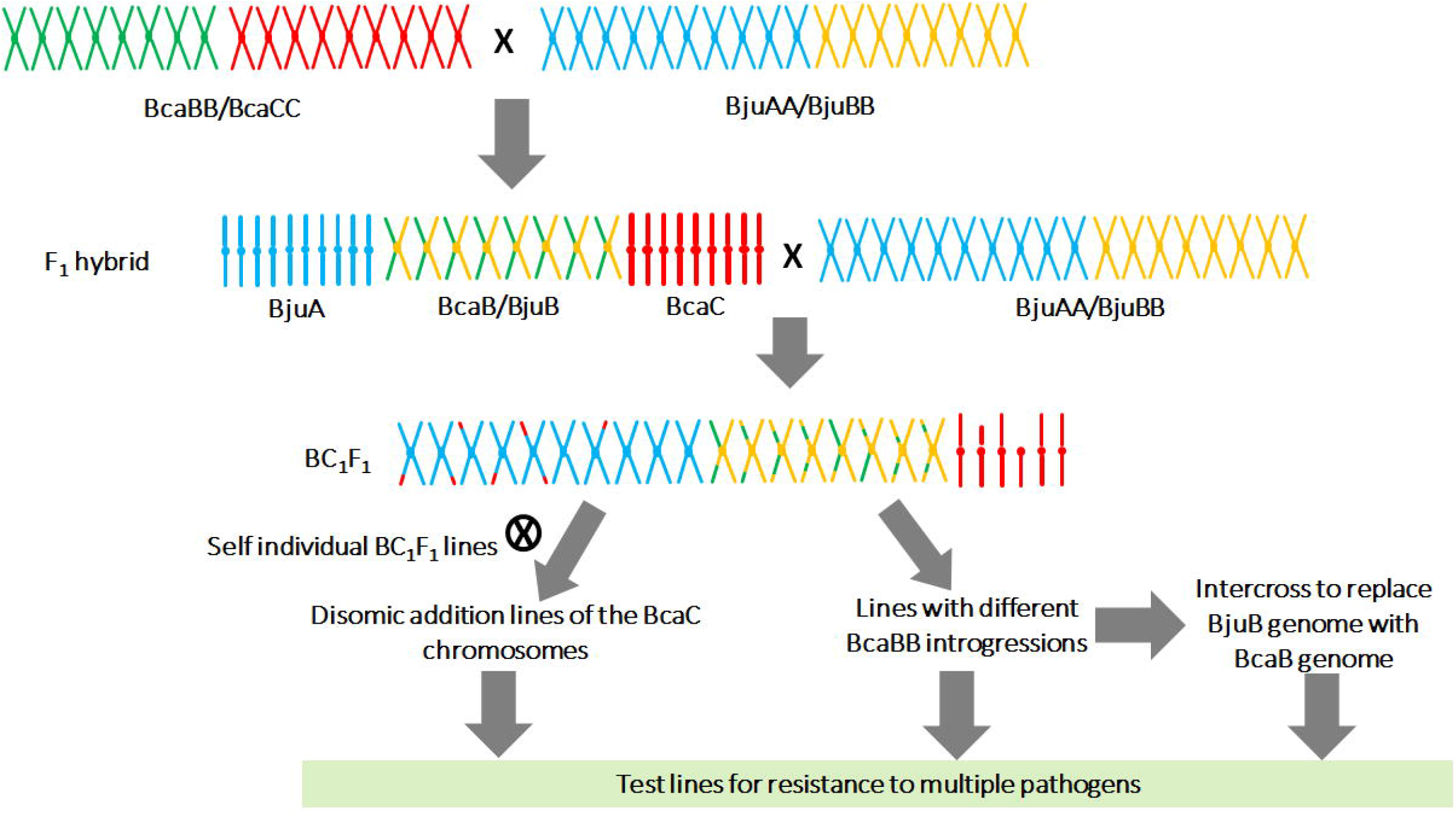
A model for genomics-assisted mapping and introgression of resistanceconferring loci in inter-specific crosses between *B. carinata* and *B. juncea*. Two independent strategies have been proposed for identifying the resistance-conferring regions in the BcaB and BcaC genomes in the background of Bju genome. In case of mapping of resistance-conferring regions from BcaC genome, generation of disomic addition lines of different BcaC chromosomes followed by phenotyping needs to be carried out. For mapping resistance-conferring regions from BcaB genome, different introgression lines of BcaB into BjuB need to be genotyped at the BC_1_F_1_ stage. The BC_1_F_1_ lines containing different BcaB introgressions can either be selfed to fix the introgressions or intercrossed to obtain a nearcomplete replacement of BjuB with BcaB genome.

We propose two independent strategies for identifying the resistance-conferring regions in the BcaB and BcaC genomes. The BcaB and BjuB subgenomes showed high recombination frequency in the interspecific crosses. For biotrophic pathogens, wherein the resistance is mostly governed by NLR genes, selective introgressions of NLR clusters of the BcaB genome in the BjuB background can be identified and the progenies tested for resistance. For necrotrophic pathogens like *S. sclerotiorum* and *A. brassicae*, the resistance is usually quantitative, wherein multiple genes contribute to resistance (Rajarammohan *et al*., 2018; Roy *et al*., 2021; Wu *et al*., 2013; Wu *et al*., 2019). In this scenario, the near-complete replacement of BjuB with BcaB could be achieved by intercrossing BC_1_F_1_ plants with different BcaB introgressions to create new genetic diversity in the *B. juncea* gene pool that may confer resistance to the necrotrophic pathogens. However, caution must be exercised to select lines with minimal changes to the BjuA chromosomes as the BjuA subgenome contributes maximally to the yield of *B. juncea* (Ramchiary *et al*., 2007; Yadava *et al*., 2012), and structural changes in the BjuA chromosomes may affect the yield.

To identify disease resistance loci in the BcaC genome, disomic addition lines for each BcaC chromosome in the *B. juncea* background would be the best strategy. The disomic addition lines of the BcaC chromosomes can be obtained from the self-pollination of the BC_1_F_1_ plants with one or more BcaC monosomic additions and assayed for disease resistance. Although previous attempts to create an allohexaploid (AABBCC) *Brassica* population resulted in the loss and duplication of A and C genome chromosomes (Mason *et al*., 2014b), it would be interesting to study the stability of the C genome disomic addition lines in the *B. juncea* (AABB) background.

*B. carinata* can also be used to broaden the genetic diversity in *B. napus*, particularly for the common C subgenome. However, structural differences between the BnaA and BcaB genomes will pose difficulties for introgressions from BcaB to BnaA.

### Importance of *B. carinata* as an oilseed & vegetable crop in Africa

*B. carinata* is projected as a biofuel crop in the developed parts of the world (Taylor *et al*., 2010). However, *B. carinata* could be an important crop to boost edible oil availability in Africa. Both *B. carinata* (grown predominantly in Africa) and *B. juncea* (grown predominantly in the Indian subcontinent) are important crops for low-input, high-output agriculture. The current yields of *B. carinata* are much lower as compared to the other allopolyploids - *B. napus* and *B. juncea*. A backcross of the initial interspecific hybrid (BBAC) to *B. carinata* may allow for the introgression of BjuA into BcaC. The current *B. juncea* cultivars are high-yielding with yield-related QTLs mapped to regions on BjuA07, BjuA08, and BjuA10 (Ramchiary *et al*., 2007; Yadava *et al*., 2012). The interspecific transfer of yield-QTL-containing regions from *B. juncea* to *B. carinata* may therefore help to improve yields of *B. carinata*.

In conclusion, with the availability of highly contiguous genome assemblies and abundant SNP markers – interspecific crosses between *B. carinata* and *B. juncea* could lead to the improvement of the two oilseed crops and benefit the small farm holders both in South Asia and Africa.

## Materials and Methods

### Plant material, genome size estimation, nanopore sequencing and genome assembly

*B. carinata* accessions HC20, HC25, NPC2, and BEC184 used for genome sequencing experiments had been maintained by self-pollination for more than five generations. For DNA isolation, seedlings were grown in a growth chamber at 8 h light, 25°C and 16 h dark, 10°C cycle. Leaves were harvested from ~15-day-old seedlings and immediately snap-frozen in liquid nitrogen. High-molecular-weight DNA was isolated from the frozen tissues by the CTAB method (Rogers and Bendich, 1994).

For Nanopore sequencing, genomic DNA libraries were prepared from ~5 μg of high molecular weight DNA using the ‘Ligation sequencing kit 1D’ following the manufacturer’s instructions (Oxford Nanopore). The quality and quantity of the DNA libraries were determined with a Nanodrop spectrophotometer. DNA libraries were sequenced on the MinION and PromethION devices using the MinION and PromethION Flow Cells R9.4.1 (Oxford Nanopore). Base-calling and quality filtering were performed using Guppy software (https://github.com/guppy).

For Illumina sequencing, ~5 μg of genomic DNA was fragmented using a Covaris ultrasonicator. Fragmented DNA was used to prepare Illumina paired-end (PE) libraries of size 150 bp for *B. carinata* accessions HC20, HC25, NPC2, and BEC184 following the manufacturer’s recommended protocol (https://illumina.com). The PE libraries were sequenced on Illumina HiSeq 2000 system at ~40x depth for HC20 and ~10x for the rest of the accessions.

A kmer frequency distribution analysis was carried out with the Illumina PE reads (kmer length – 25) to estimate the genome size using Jellyfish v2.2.6 (Marçais and Kingsford, 2011). Raw Nanopore reads were assembled into contigs using the Canu assembler V1.6 (Koren *et al*., 2017) with the parameters ‘minRead length’ and ‘minOverlap length’ set at values of 1000 bp. The data was assembled on a Lenovo SR950 server with 392 cores and 1 TB RAM. The paired-end reads obtained with Illumina sequencing were mapped on the assembled Nanopore contigs using BWA-MEM (Li and Durbin, 2009), followed by error correction with the Pilon tool (Walker *et al*., 2014) in five iterative cycles. After each of the Pilon cycles, the accuracy of the corrected genome was ascertained using the MUMmer tool (Kurtz *et al*., 2004).

### DLS-based optical mapping

Optical mapping was performed following the protocols suggested by the manufacturer (Bionano Genomics; https://bionanogenomics.com/). Leaf tissues from seven-day-old seedlings were harvested and transferred to an ice-cold fixing solution. Nuclei were isolated using the ‘rotor-stator’ protocol (Bionano Genomics, document no: 30228) and purified on a sucrose density gradient. The nuclei were embedded in 0.5 % w/v agarose and treated with Proteinase-K (Qiagen) for 2 h. Mapping was carried out using the DLS (Direct Label and Stain) system. The DNA was labeled with the DLE-1 enzyme using the ‘Bionano Prep DLS kit’. Mapping data were obtained from the labeled libraries on the Saphyr system (Bionano) using one lane for each library. Optical maps and hybrid assemblies were generated using the Bionano Access software.

### Genotyping and genetic map construction

A genetic map was constructed using 181 F_1_DH lines derived from a cross of HC20 and BEC184. Genotyping of the 181 F_1_DH lines was carried out using the Illumina Infinium™ Brassica 90k Array (Clarke *et al*., 2016; Scheben *et al*., 2019). To generate additional markers, 94 F_1_DH plants were genotyped using genotyping by sequencing (GBS) with the SphI-MlucI restriction enzyme combination (Fu *et al*., 2016). Genetic mapping was carried out in the R statistical computing environment (R Core Team 2021; RStudio Team 2021) using the ASMap package (Taylor and Butler, 2017). The R/ASMap package uses the R/qtl (Broman *et al*., 2003) input format and is based on the MSTmap algorithm (Wu *et al*., 2008). The “all-in-one” function mstmap was implemented in R/ASMap with the following parameters: distance function ‘Kosambi’, a minimum cut-off p-value of 1e-6 and a missing threshold of 0.3. The physical marker positions were used to combine the minor unassigned linkage groups with >7 markers with the known linkage groups.

### Comparative genome analysis of the U’s triangle species

Six genome assemblies (*B. rapa* Chiifu v3.0, *B. nigra* Sangam, *B. oleracea* HDEM, *B. juncea* Varuna, *B. napus* darmor-bzh v10, and *B. carinata* HC20) representing the six species of the U’s triangle were aligned using LAST and filtered using the cscore parameter (cscore > 0.99) to obtain the reciprocal best hit for each gene. The homologous regions were identified and aligned as anchors using the genes. To calculate the Ks values of the collinear gene pairs, the CDS sequences of the gene pairs were aligned using MAFFT (Katoh *et al*., 2002) and the Ka, Ks, and Ka/Ks values were calculated using the NG model in the KaKs_Calculator program (Zhang *et al*., 2006). Further, the 23 long-read Brassica genomes (Table S16) were compared using the MCScan (Python version) (Wang *et al*., 2012) by aligning the respective subgenomes of the allopolyploids with the diploid genomes. Macrosynteny plots were then generated to visualize and detect any significant structural rearrangements.

### Phylogenomic analysis of the U’s triangle species

To determine the phylogenetic relationship among the six species of the U’s triangle, the protein sequences of the 23 genomes were obtained from the data repositories mentioned in their respective publications (Table S16). The protein sequences of the allopolyploids were separated according to their constituent subgenomes and *A. thaliana* was used as the outlier. Gene family clusters were identified using OrthoFinder V 2.2.7 (inflation value: 1.5) (Emms and Kelly, 2019). The clusters were further analyzed with Mirlo (https://github.com/mthon/mirlo) to identify phylogenetically informative single-copy gene families. These families were then concatenated into one large alignment and trimmed using Gblocks V 0.91.1 (Talavera and Castresana, 2007) to remove ambiguous regions. The concatenated alignment was then used to construct a phylogenetic tree by FastME V 2.1.6.1 (Lefort *et al*., 2015) with bootstrap support values (500 replicates) calculated by Booster V 0.3.1 (Lemoine *et al*., 2018).

### NLR prediction and annotation

NLRs were identified using the RGAugury pipeline (Li *et al*., 2016), which identifies and classifies three types of resistance genes, NLR, RLK, and RLP. The protein and DNA sequences of the six Brassica species (*B. rapa* Chiifu v3.0, *B. nigra* Sangam, *B. oleracea* HDEM, *B. juncea* Varuna, *B. napus* darmor-bzh v10, and *B. carinata* HC20) were used as input and the pipeline was run using the default parameters. Additionally, the NLR-Parser tool (Steuernagel *et al*., 2015) was also used to independently annotate the NLRs in the six genomes. The predicted NLR genes from both pipelines were manually inspected and curated to obtain the final NLR repertoire of each species. The RPW8 domain containing NLR proteins were checked manually using NCBI Conserved Domain Database (CDD) tool (https://www.ncbi.nlm.nih.gov/Structure/cdd/wrpsb.cgi) to confirm their classification.

### Generation of *B. carinata* x *B. juncea* interspecific hybrids and SNP genotyping of the progenies

For developing *B. carinata* x *B. juncea* interspecific hybrids, *B. juncea* Varuna was crossed with *B. carinata* accessions HC-20, HC-25 and NPC-2 with Varuna as the female parent. The details of the crosses and embryo rescue are given in Figure S19. A total of 8, 4, and 17 F_1_ plants of the VHC1, VHC2, and VNP crosses, respectively, were transferred to the field and backcrossed with Varuna as the recurrent parent.

Genomic DNAs were isolated from young leaf tissues of the field-grown BC_1_F_1_ and BC_2_F_1_ plants using the CTAB method. Genotyping of the BC_1_F_1_ progenies was performed using SNPs obtained from the Illumina Infinium™ Brassica 90K Array (Clarke *et al*., 2016; Scheben *et al*., 2019). Hybridization was performed according to the manufacturer’s instructions for the samples, and the genotype data were exported using Genome Studio v 2.0.4 software. A total of 50,018, 49,434 and 41,500 SNPs were obtained for VNP, VHC2, and VHC1 BC_1_F_1_ plants (Table S22). We filtered out heterozygous SNPs that hybridized with both the A and C genomes. BjuA- and BcaC-specific SNPs that hybridized only to the corresponding parental DNA were identified and physically mapped on the Varuna and HC20 genome assemblies. Information on the BjuA- and BcaC-specific SNPs in the three different crosses are described in Table S22. For the B genome, SNPs that were polymorphic between the BjuB and the BcaB genome in each cross were used. Karyotypes were generated using custom R scripts.

## Acknowledgments

The research was supported by the Department of Biotechnology (DBT), Government of India, under the projects ‘DBT-UDSC partnership Centre on Genetic Manipulation of Brassicas’ (grant number – BT/01/NDDB/UDSC/2016), and ‘Centre of Excellence on Genome Mapping and Molecular Breeding of Brassicas’ (grant number - BT/01/COE/08/06-II). SR is supported by DST-INSPIRE Faculty fellowship grant (IFA18-LSPA118); DP is supported by a SERB - National Science Chair.

## Authors contributions

KP, AKP, JK, and DP conceived the project. KP and SR carried out the sequencing, genome assembly, transcriptome, SNP genotyping, and comparative genome analyses. SKY, RV, AM, AKP developed DH lines and carried out genetic mapping. SS, RV, AM, and VG made the interspecific hybrids and the subsequent backcrosses. SM helped with the transcriptome work. SR, KP, and DP wrote the early drafts. All authors read and approved the final version of the manuscript.

## Code availability

Codes used in the manuscript are available on request.

## Data availability statement

All the raw data generated in the manuscript have been deposited in the Indian Nucleotide Data Archive (INDA) under the study accession: INRP000031. The data is also mirrored at the INSDC under the BioProject ID: PRJEB55587. The *B. nigra* Sangam genome assembly from which the LG nomenclature of *B. carinata* B chromosomes has been derived is deposited under the NCBI BioProject ID: PRJNA642332. The *B. juncea* Varuna updated genome assembly is deposited under the NCBI BioProject ID: PRJEB51129.

## Supplementary Information

### Supplementary Tables

Table S1: *B. carinata* HC20 raw sequencing data obtained from Oxford Nanopore minION and PromethION

Table S2: *B. carinata* HC20 Illumina raw sequencing data

Table S3: Mapping data generated at each step of the hierarchical optical mapping analysis of *B. carinata* HC20 genome

Table S4: Characteristics of the linkage map of *B. carinata* developed using HC20 x BEC184 F1DH mapping population

Table S5: Position and number of the assembled scaffolds and contigs on each of the seventeen pseudochromosomes of *B. carinata*

Table S6: Nomenclature of the BcaB chromosomes of *B. carinata*

Table S7: BUSCO statistics of the *B. carinata* HC20 assembly

Table S8: Repeat summary of *B. carinata* HC20 genome

Table S9: Gene positions and block arrangement in the *B. carinata* HC20 genome

Table S10: List of flowering time genes in the representative genome assemblies of the U’s triangle species along with their chromosomal positions

Table S11: List of glucosinolate biosynthesis and catabolism genes in the representative genome assemblies of the U’s triangle species along with their chromosomal positions

Table S12: List of transcription factors in the representative genome assemblies of the U’s triangle species along with their chromosomal positions

Table S13: *B. carinata* transcriptome raw data statistics

Table S14: List of DEGs in various tissues of *B. carinata*

Table S15: List of tissue-specific genes expressed in the leaf, root, and seeds of *B. carinata*

Table S16: Summary of long-read genomes of *Brassica* species used for comparative analyses

Table S17: Summary of genes assigned to orthogroups in the comparison of U’s Triangle species

Table S18: NLR genes in the *B. carinata* HC20 genome and their classification

Table S19: Sensor-Executor NLR gene pairs in the *B. carinata* HC20 genome

Table S20: Summary of the comparative analysis of NLR genes in the U’s Triangle

Table S21: Raw read statistics of resequencing of *B. carinata* accessions NPC2, BEC184, and HC25

Table S22: Summary of SNP markers obtained from the Illumina 90K Brassica genotyping array for the interspecific crosses between *B. carinata* and *B. juncea*

Table S23: Chromosome counts of *B. carinata* x *B. juncea* interspecific hybrids

Table S24: Heterozygosity status of the B genome of BC_1_F_1_ plants from *B. carinata* x *B. juncea* crosses

Table S25: Heterozygosity status of the B genome of BC_2_F_1_ plants from *B. carinata* x *B. juncea* crosses

### Supplementary Figures

Figure S1: Genome size estimation of *B. carinata* HC20 using k-mers

Figure S2: Quality improvement of the HC20 assembly with each step of polishing (number of SNPs/indels)

Figure S3: Circos plot denoting the position of flowering time genes in the six Brassica species.

Figure S4: Circos plot denoting the position of glucosinolate genes in the six Brassica species.

Figure S5: Comparative analysis of 58 transcription factor families across the six *Brassica* species

Figure S6: Pearson correlation plot of the samples used in the transcriptome analysis

Figure S7: Leaf-specific expression profiles of genes in the *B. carinata* HC20 accession

Figure S8: Root-specific expression profiles of genes in the *B. carinata* HC20 accession

Figure S9: Seed-specific expression profiles of genes in the *B. carinata* HC20 accession

Figure S10: Whole-genome comparisons of AA subgenomes of *B. juncea* and *B. napus* long-read genomes with *B. rapa*.

Figure S11: Whole-genome comparisons of BB subgenomes of *B. juncea* and *B. carinata* long-read genomes with *B. nigra*.

Figure S12: Whole-genome comparisons of CC subgenomes of *B. carinata* and *B. napus* long-read genomes with *B. oleracea*.

Figure S13: Boxplots of Ks values of *B. juncea, B. carinata, B. napus* subgenomes with the diploid genomes *B. rapa*, *B. nigra* and *B. oleracea*.

Figure S14: Phylogenetic analysis of RPW8 domain-containing proteins of *B. carinata*.

Figure S15: Maximum-likelihood phylogenetic trees of ‘sensor–executor’ paired NLR genes identified in HC20 genome.

Figure S16: Position of NLR genes along the A chromosomes of *B. rapa, B. juncea*, and *B. napus*.

Figure S17: Position of NLR genes along the B chromosomes of *B. nigra, B. juncea*, and *B. carinata*.

Figure S18: Position of NLR genes along the C chromosomes of *B. oleracea, B. carinata*, and *B. napus*.

Figure S19: Schema of the interspecific crosses between *B. carinata* x *B. juncea* across the two generations

Figure S20: Chromosomal makeup of BC_2_F_1_ plants of the J29 BC_1_F_1_ line

Figure S21: Chromosomal makeup of BC_2_F_1_ plants of the J40 BC_1_F_1_ line

Figure S22: Chromosomal makeup of BC_2_F_1_ plants of the J52 BC_1_F_1_ line

